# Construct a variable-length fragment library for *de novo* protein structure prediction

**DOI:** 10.1101/2022.01.03.474755

**Authors:** Qiongqiong Feng, Minghua Hou, Jun Liu, Kailong Zhao, Guijun Zhang

## Abstract

Although remarkable achievements, such as AlphaFold2, have been made in end-to-end structure prediction, fragment libraries remain essential for *de novo* protein structure prediction, which can help explore and understand the protein-folding mechanism. In this work, we developed a variable-length fragment library (VFlib). In VFlib, a master structure database was first constructed from the Protein Data Bank through sequence clustering. The Hidden Markov Model (HMM) profile of each protein in the master structure database was generated by HHsuite, and the secondary structure of each protein was calculated by DSSP. For the query sequence, the HMM-profile was first constructed. Then, variable-length fragments were retrieved from the master structure database through dynamically variable-length profile-profile comparison. A complete method for chopping the query HMM-profile during this process was proposed to obtain fragments with increased diversity. Finally, secondary structure information was used to further screen the retrieved fragments to generate the final fragment library of specific query sequence. The experimental results obtained with a set of 120 nonredundant proteins showed that the global precision and coverage of the fragment library generated by VFlib were 55.04% and 94.95% at the RMSD cutoff of 1.5 Å, respectively. Compared to the benchmark method of NNMake, the global precision of our fragment library had increased by 62.89% with equivalent coverage. Furthermore, the fragments generated by VFlib and NNMake were used to predict structure models through fragment assembly. Controlled experimental results demonstrated that the average TM-score of VFlib was 16.00% higher than that of NNMake.

## 1 Introduction

Computationally predicted three-dimensional structure of protein molecules has demonstrated the usefulness in many areas of biomedicine, ranging from approximate family assignments to precise drug screening [1]. Prediction based on co-evolutionary information by using deep learning has considerably advanced protein structure prediction [2–4]. However, current deep learning methods, including the end-to-end structure prediction methods [5, 6], have limited effectiveness in the exploration and understanding of the protein-folding mechanism that is one of the central problems in biochemistry and essential for the study of protein misfolding diseases [7–9].

Fragment-based *de novo* protein structure prediction methods are important for understanding the protein-folding mechanism, including Rosetta [10], QUARK [11, 12], CGLFold [13], and so on. In Rosetta, short fragments of known proteins are assembled by a Monte Carlo strategy to yield native-like protein conformations. For each structure prediction, many short simulations starting from different random seeds are carried out to generate an ensemble of decoy structures that have both favorable local interactions and protein-like global properties. This decoy set is then clustered by structural similarity to identify the broadest free energy minima. In QUARK, the algorithm starts from position-specific fragment structure generation. Replica-exchange Monte Carlo (REMC) simulations are performed to assemble the fragments into full-length models under a composite physics- and knowledge-based potential. In CGLFold, fragment recombination and fragment assembly are used to implement large-scale conformation space search to form a topology. Then, a loop-specific perturbation model is constructed to further improve the conformation formed in the global phase. However, the performances of fragment assembly-based structure prediction methods are well known to depend on the quality of fragment libraries [14].

NNMake, the fragment library generation algorithm used in Rosetta, searches for 9- and 3-residue fragments from a nonredundant structure database [15]. The fragments are scored based on primary sequences as well as predicted secondary structures and torsion angles, and the top 200 candidates are selected for each position of the target protein. Despite the success of Rosetta in many applications, the protocol of finding a fixed number of fragments with a fixed length at each protein position may have limited the quality of fragments identified by NNMake [14, 15]. HHfrag [16] is a dynamic HMM-based fragment search method built on the profile-profile comparison tool HHpred [17]. HHfrag obtains fragments with lengths from 6 to 21 residues using dynamic HMM by searching PDBselect25 [18]. However, sequence homologs were not excluded from the HHfrag results, which may inflate the method’s precision. And for some target positions, HHfrag does not output any fragments, which may cause difficulties during the modelling step [19]. Flib [19] extracts fragments from a template database using a combination of random and exhaustive approaches [20]. A library containing the top-3000 fragments per position is compiled by using the secondary structure score and the Ramachandran specific sequence score (LIB3000). LIB3000 is then sorted according to the torsion angle score and the top-20 fragments per position are selected to construct the final library. LRfraglib [14] first built logistic regression models to evaluate the quality of fragments of 7-10 residues, using the similarity scores of primary sequences, physicochemical properties and secondary structures, and then developed a multistage fragment-selection protocol with these models [14]. Previous studies have demonstrated that fragments are essential for fragment assembly-based methods. However, current fragment libraries remain inadequate, and the accuracy of prediction models generated by fragment assembly will be difficult to reach expectations when fragment libraries lack fragments with near-native fragment structures [21, 22]. Designing a variable-length fragment library with increased completeness is crucial for improving the accuracy of prediction models.

In this work, a master database was first constructed, and then variable-length fragments were retrieved from the master structure database through dynamically variable-length profile-profile comparison. Finally, secondary structure information was utilized to further screen the retrieved fragments to generate the final fragment library of the query sequence. In addition, in fragment assembly-based protein structure prediction, each fragment was assigned a different selection probability on the basis of the predicted distance to improve the efficiency of selecting near-native structure fragments. The results of controlled experimental compared with NNMake show that the global precision of VFlib had increased by 62.89% with equivalent coverage, and VFlib achieves a higher TM-score for 96 out of 120 target proteins.

## 2 Materials and Methods

The pipeline of VFlib is presented in Figure 1. First, the master structure database is constructed by using MMseqs2 [23, 24], HHsuite [25], and DSSP [26, 27] on the basis of the latest Protein Data Bank (PDB, 2021-08). Then, the HMM-profile of query sequence is constructed and chopped into a nested set of segments (3-21). Profile-profile comparison-based fragment extraction and secondary structure information-based fragment selection strategies are used to construct the variable-length fragment library. Finally, in fragment assembly-based protein structure prediction, each fragment is assigned a different selection probability based on the predicted distance, and the final prediction model is generated by probabilistically biased fragment assembly. The distance evaluation function is used to improve the sampling efficiency.

**Figure 1.**
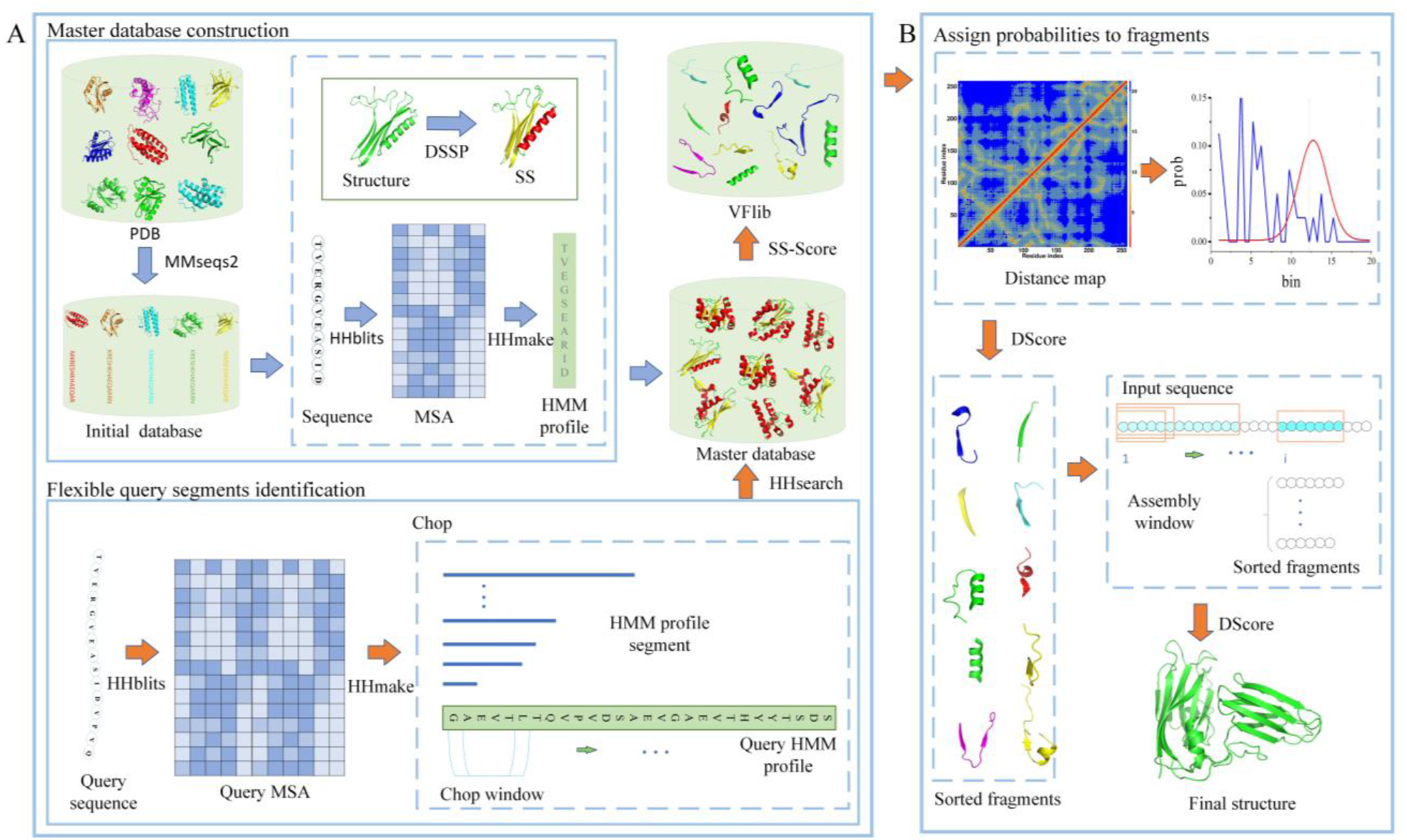
**(A)** The pipeline of VFlib. The construction of the master database (shown by the blue arrow), including the clustering of sequences in PDB, the generation of the HMM-profile, and the calculation of secondary structure distribution. The generation of the variable-length fragment library is shown by the orange arrow. First, the HMM-profile of the query sequence is generated by HHsuite and chopped into a nested set of segments (3-21). Then, the fragments are obtained by profile-profile comparison. Finally, the secondary structure information is used to further screen the fragments to obtain final variable-length fragment library of specific query sequence. **(B)** Process of fragment assembly-based protein structure prediction (shown by the orange arrow), including the assignment of different selection probabilities to fragments and the using of the distance function to guide assembly.

### 2.1 Master structure database construction

The master structure database is generated from PDB for searching the fragments of the query sequence. First, sequences and structures are obtained from PDB. Then, the initial structure library is generated by MMseqs2 with a coverage threshold of 25% and a paired sequence identity cutoff of 0.3. Finally, in the initial structure library, 11 648 proteins are retained after excluding the proteins with sequence lengths out of the range of 50-500 residues or those with corresponding PDB structures with resolutions exceeding 2.5 Å. The multiple sequence alignment (MSA) [28] for each sequence in the initial structure library is generated by HHblits [29] with a coverage threshold of 50%, a pairwise sequence identity cutoff of 0.99 and a E-value threshold of 0.001 against Uniclust30_2018_08 [30] with three iterations. Then, HHmake [25] is used to build the HMM-profile of each sequence in the initial structure library. For each structure in the initial structure library, the secondary structure assignment is obtained by DSSP [26, 27]. The master database, which includes HMM-profiles and secondary structure assignments, is thus completed.

### 2.2 Flexible query segment identification

Given that fragments of fixed lengths may be unideal, a strategy for the identification of flexible query segments is proposed to obtain fragments with increased diversity. First, all redundant proteins with pairwise sequence identity > 30% between the master library and the query are removed by CD-HIT [31] to exclude the homologs of the query protein from the master libraries to simulate a *de novo* protein structure prediction scenario correctly [32]. Then, the HMM-profile of the query sequence is generated by a process that is the same as the HMM-profile generation of the protein sequence in the master database. Finally, the query HMM-profile is chopped into variable segments with different lengths (3-21) from the insertion position which is selected starting from the first residue of the sequence. The chopping process continues until the index of the selected residue plus the length of the fragment HMM-profile exceeds the length of the query sequence. For example, when we select the *i* th residue in the query sequence, the HMM-profile of the 3 segments at the insertion position is from the *i* th residue to the *i*+2 th residue of the query HMM-profile and that of 21 segments at the insertion position is from the *i* th residue to the *i*+20 th residue.

### 2.3 Fragment extraction

After chopping the query HMM-profile, each insertion position in the query sequence contains the HMM-profiles of segments with variable length. The candidate fragments at each insertion position are obtained by using the HHsearch [33] to search the master database in accordance with the HMM-profiles of the variable segments at the insertion position. In the process, the searched fragments are placed in the candidate fragment library if the matching probability is greater than 20%. If the number of candidate fragments is less than 50, the remaining matching fragments are selected as the candidate fragments in accordance with the matching probability until the number of candidate fragments reaches 50. Each insertion position in the query sequence containing all of the HMM-profile of segments with variable lengths (3-21) has at least 950 candidate fragments.

### 2.4 Fragment selection

The length of different candidate fragments is not fixed, which may lead to the situation that a shorter candidate fragment is a part of a longer candidate fragment. In addition, some candidate fragments may have gaps. Fragment selection is designed to improve the quality of the fragment library in response to the above-mentioned situations. First, fragments with gaps are removed from the candidate fragment library, taking into account the redundancy between the fragments and the effectiveness of the fragments. Then, the candidate fragments that were searched for the same insertion position are judged whether they are from the same region of the same protein in the master database. If multiple candidate fragments originate from the same region, only the fragment with the best matching probability was retained. Finally, the retained fragments are scored on the basis of the secondary structure information of the query sequence, and the fragments below the scoring threshold are removed. The probability that each residue in the query sequence is predicted to be an α-helix, β-sheet, or loop acquired from the PSIPRED [34] is used as the criterion for the secondary structure of residues. The native secondary structure information of all candidate fragments is obtained from the corresponding proteins in the master database. The candidate fragments derived in the previous steps are scored in accordance with the 3 × 3 matrix shown in Supplementary Table S1 and normalized to obtain the score of each residue. Candidate fragments with secondary structure score (SS_score) higher than the set threshold was selected to generate the final fragment library of specific query sequence. The details about fragment selection are shown in Supplementary Figure S1.

### 2.5 Assignment of probabilities to fragments

The method of fragment assembly generally involves the random selection of fragments such that each fragment at the insertion position has the same probability of being selected [22, 35, 36]. However, the strategy of random fragment selection does not consider that near-native structure fragments should be preferentially selected, which may result in low assembly efficiency. Therefore, a strategy for assigning different selected probabilities to fragments on the basis of distance information is proposed. The predicted distance information of the query sequence is obtained from the trRosetta [4], and the variable-length fragments generated by VFlib are assigned probabilities on the basis of the predicted distance. In accordance with the fragment length of the insertion position, the predicted distance information of the corresponding residue pair is selected to generate the probability distribution curve of the distance between the residue pairs via Gaussian fitting. At the same time, the distance of the fragment is calculated and compared with the corresponding distance in the distance probability distribution curve to determine the selected probability. Finally, the probabilities of all fragments at the insertion position are normalized to increase the convenience and reliability of selecting fragments for assembly.

### 2.6 Distance-guided fragment assembly

The accurate potential energy function is very important for the sampling efficiency of the fragment assembly. An energy function based on the distance information is designed to select the conformation. First, the distance information of the query sequence obtained by the trRosetta [4] is filtered to focus on critical information. For example, the residue pair distances with the highest probability for the first bin (that is, the bin whose predicted distance exceeds 2-20 Å) are removed. Then, Gaussian fitting is performed on the distance distribution of the remaining residue pairs to obtain the distance mean and variance. Finally, the distance mean and variance of these residue pairs are used to build the potential energy function in accordance with Equation 1.

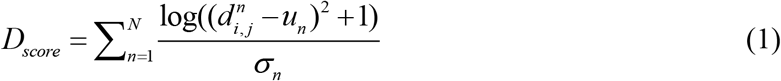

where *N* is the number of effective inter-residue distances; *i* and *j* are the residue indexes of the *n* th distance; *d_i,j_* is the real distance between *C_β_* (*C_α_* for glycine) of residues *i* and *j* in the evaluated conformation; and *u_n_* and *σ_n_* are the mean and standard deviation obtained by the Gaussian fitting of the *n* th inter-residue distance distribution, respectively.

In the process of fragment assembly, fragments are selected on the basis of the distance information to replace the residues in the corresponding region at the randomly selected insertion position of the query sequence. The Metropolis criterion based on the distance energy function is used to determine whether the updated conformation is accepted.

## 3 Results

In this work, the quality of the fragment library generated by the VFlib and NNMake was evaluated on 120 nonredundant benchmark proteins. The influence of secondary structures on the quality of the fragment library was discussed. At the same time, fragments were used for fragment assembly with Rosetta’s ClassicAbinitio protocol [10] to further evaluate the quality of fragment library.

### 3.1 Evaluation of the fragment library

Two commonly utilized metrics of global precision [37] and coverage [38] are used to evaluate the quality of fragment libraries generated by VFlib and NNMake on 120 nonredundant benchmark proteins. Precision is the proportion of near-native fragments, and coverage is the proportion of positions that are covered by at least one near-native fragment; near-native fragments are fragments that are structurally close to the native fragment with a certain RMSD cutoff value [14]. Considering that RMSD strongly depends on fragment length and the fragments generated by VFlib have variable lengths, we applied normalized RMSD [39]. Specifically, the RMSD values of the 6-21 fragments were normalized to that of 9-residue fragments, and the RMSD values of the 3-5 fragments were normalized to that of 3-residue fragments in accordance with Equation 2.

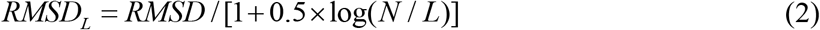

where *N* is the number of residues in the fragment, and *RMSD_L_* is the normalized value for *L*-residue fragments.

Figure 2 shows a comparison of the quality of the fragments generated by VFlib (6-21 fragments) and NNMake (9 fragments). The global precision of the fragments generated by VFlib has improved at each RMSD cutoff value (0-2 Å), and the coverage is equivalent. A fragment is considered as a true hit or true positive if the RMSD to the native structure is below a threshold value (typically 1.5 Å) [16]. At the RMSD cutoff of 1.5 Å, the precision of the fragment library generated by VFlib (55.04%) is 62.89% higher than that of NNMake (33.79%), and the coverage of VFlib (94.95%) has reduced by 2.41% compared with that of NNMake (97.29%). However, fragment assembly with various variable-length fragments can compensate for the loss of coverage. The average precision and average coverage of VFlib (6-21 fragments) and NNMake (9 fragments) at each RMSD cutoff is show in Supplementary Table S2, and the detailed result of each protein at each RMSD cutoff is shown in Supplementary Table S3.

**Figure 2.**
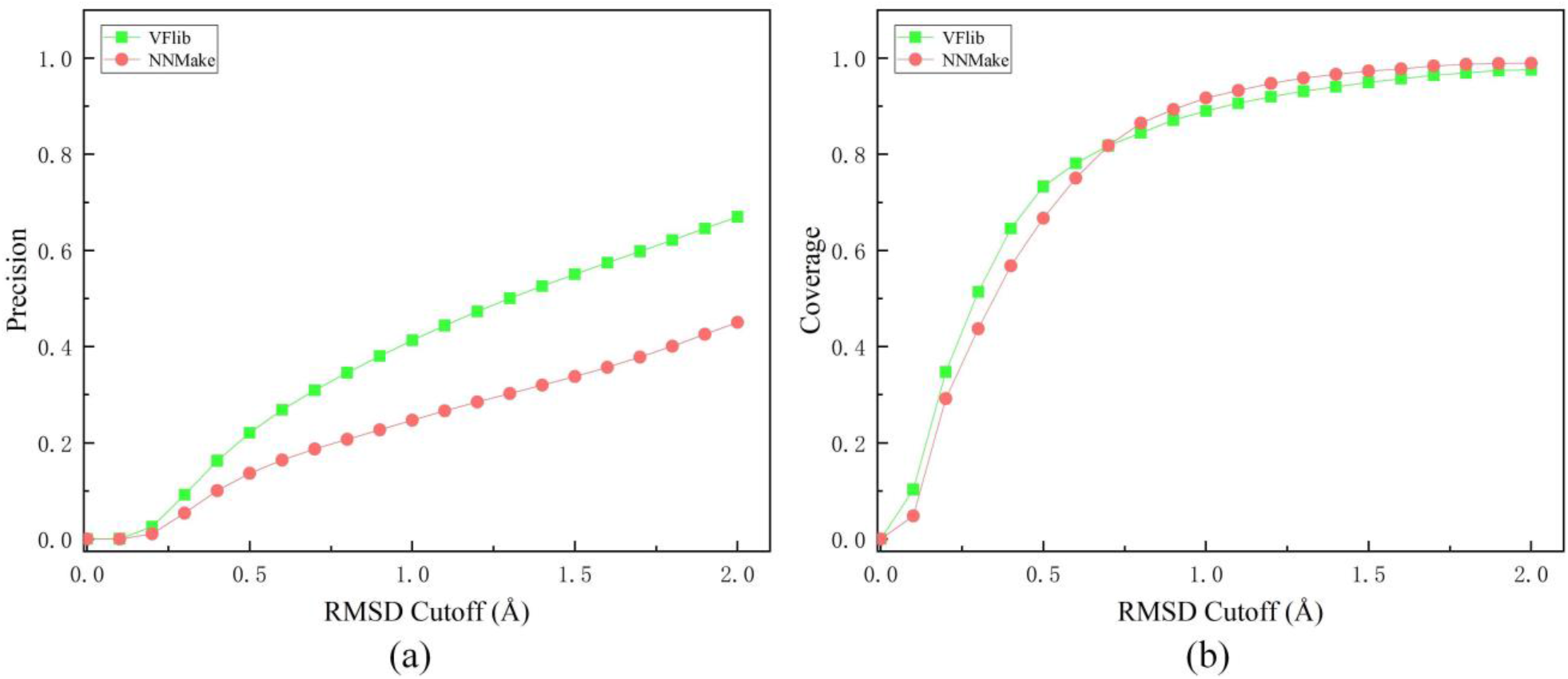
Performance of fragment libraries generated by VFlib and NNMake in the test set. VFlib and NNMake are presented by filled circles and filled squares. (a) the precision of VFlib and NNMake with different RMSD cutoff. (b) the coverage of VFlib and NNMake with different RMSD cutoff.

In addition, the precision and coverage of the fragments generated by VFlib (3-5 fragments) and NNMake (3 fragments) are high and basically similar at each RMSD cutoff value likely because of the short length of the three fragments. The average precision and average coverage of VFlib (3-5 fragments) and NNMake (3 fragments) at each RMSD cutoff is show in Supplementary Table S4 and Figure S2, and the detailed result of each protein at each RMSD cutoff is shown in Supplementary Table S5. In summary, the experimental results show that the quality of fragment library generated by VFlib is better than that of NNMake.

### 3.2 Effect of secondary structure on the fragment library

In VFlib, secondary structure information is used to further screen the fragments and improve the quality of the fragment library. As shown in Figure 3, the precision and coverage of the fragment library are different at each secondary structure score (SS_score) thresholds. With the SS_score threshold of the same RMSD cutoff value increases, the precision of VFlib improves whereas coverage deteriorates. It may be because that the low-quality fragments are continuously eliminated as the strictness of the criteria for fragment selection based on secondary structures increases. In addition, the predicted secondary structure information of some residues may be inaccurate, which may cause some high-quality fragments at the insertion position to be discarded. Therefore, it is necessary to choose a suitable SS_score of fragment selection to better assist the fragment assembly. As shown in Supplementary Table S6, if the SS_score threshold is <2.0, the coverage of the fragment library generated by VFlib is greater than 95%. The detailed results of average precision and average coverage at different SS_score thresholds and RMSD cutoff are shown in Supplementary Table S7. In order to verify further whether the selected SS_score is reasonable, the fragments obtained under different selection criteria were subjected to the same fragment assembly experiment on the same 120 nonredundant benchmark proteins. And the RMSD and TM-score were used to evaluate the prediction model’s performance [40]. Controlled experimental results show that the best prediction model was obtained when the SS_score is 2.0 (shown in Figure 4). The detailed results of average RMSD and average TM-score of each protein at different SS_score thresholds are shown in Supplementary Table S8.

**Figure 3.**
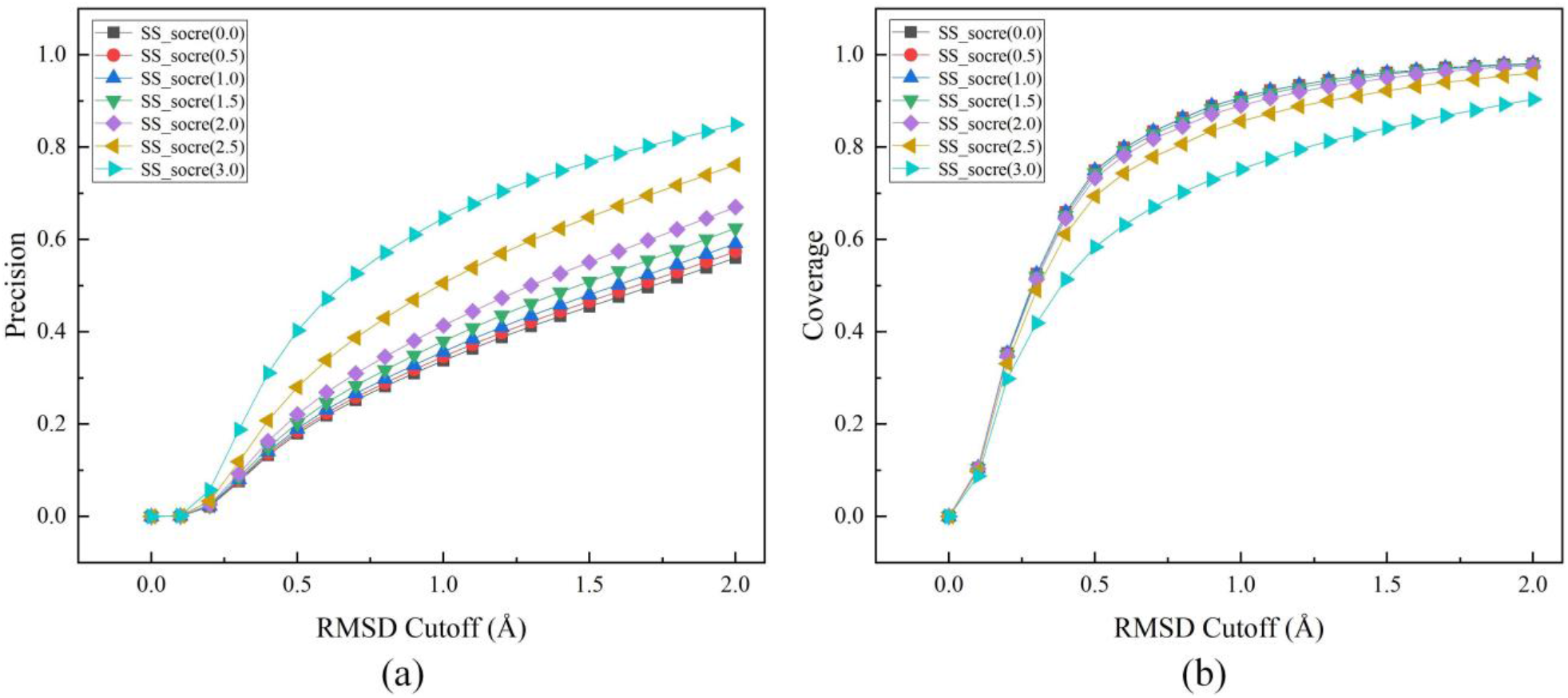
Performance of fragment libraries generated by VFlib with different SS_score thresholds. (a) the precision of fragment libraries with different RMSD cutoff. (b) the coverage of fragment libraries with different RMSD cutoff.

**Figure 4.**
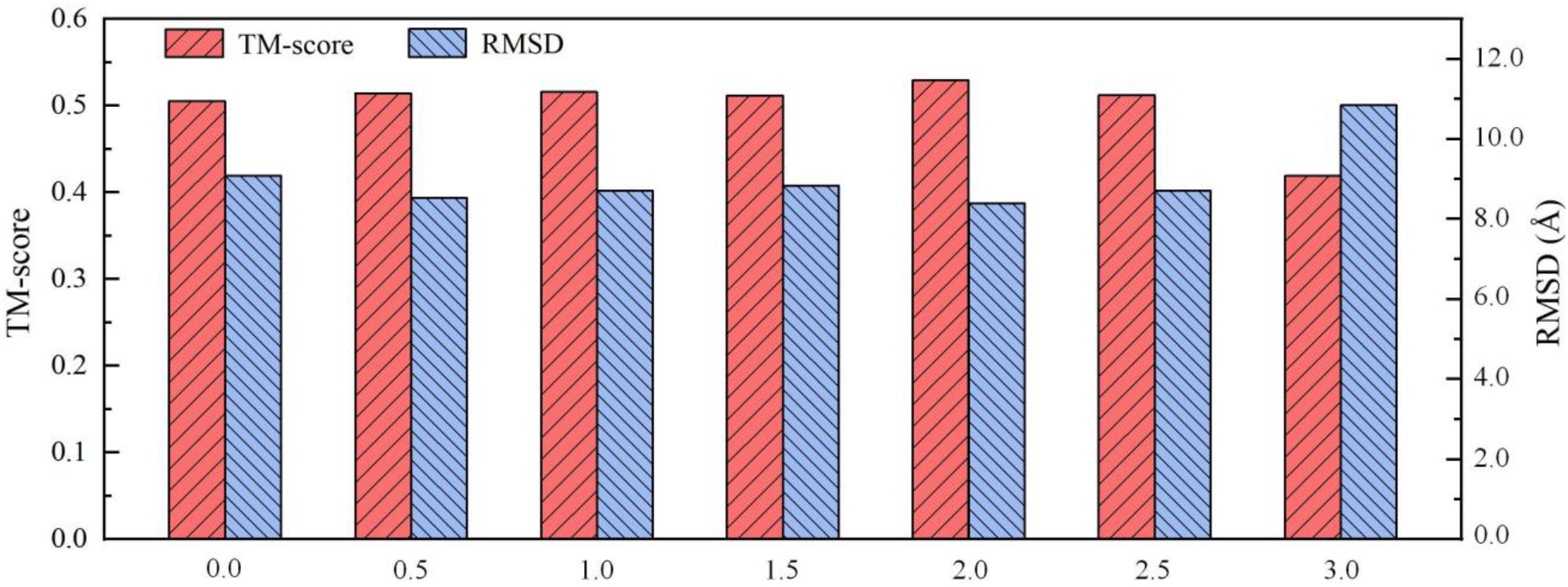
The average TM-scores and RMSD of prediction models at different SS_score thresholds. The x coordinate represents secondary structure score (SS_score).

### 3.3 Performance in protein structure sampling

In order to further evaluate the quality of the fragment library, fragments generated by VFlib and NNMake were used to by fragment assembly of Rosetta’s ClassicAbinitio protocol [10], and various experiments were conducted on the 120 benchmark proteins. In accordance with the distance information, different probabilities were assigned to the fragments generated by NNMake and VFlib. In fragment assembly-based protein structure prediction, we did not utilize the fragment assembly strategy in the Rosetta protocol because the weights in the scoring functions and the parameters in simulations were optimized for fragments of fixed lengths (3 and 9 residues), whereas VFlib uses variable fragment lengths [14]. Thus, we developed a simple *de novo* protein structure sampling program. For a given sequence and a fragment library, this program initializes a random conformation and substitutes the contiguous residue segments of the query sequence with candidate fragments in the library by using the Monte Carlo strategy. We applied an energy function that is composed of the distance information between residue pairs. The processes of fragment assembly, except for the fragment library, were the same to ensure the objectivity of the evaluation. Controlled experimental results on the same 120 nonredundant benchmark proteins are presented in Table 1. The average TM-score of the prediction models generated by using VFlib (0.58) is 16.00% higher than that of NNMake (0.50), and the average RMSD of the prediction models generated by using VFlib (7.48 Å) has reduced by 18.16% compared with that of NNMake (9.14 Å). Our method obtains 29 prediction models with TM-score >0.7, whereas NNMake obtains 10 prediction models with TM-score >0.7. Controlled experimental results in Figure 5 (a) illustrate that VFlib achieves a higher TM-score than NNMake for 96 out of 120 nonredundant target proteins.

**Figure 5.**
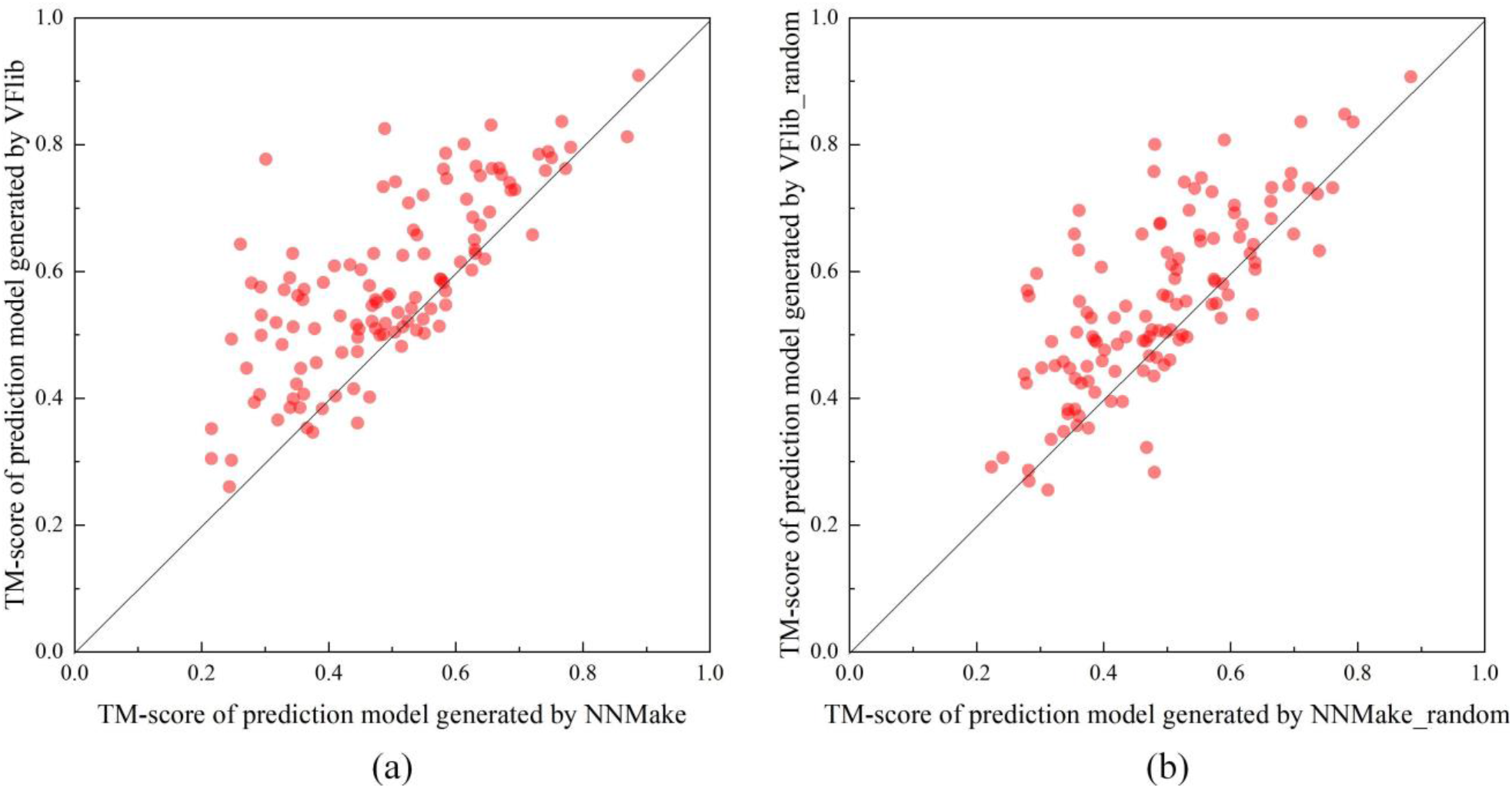
(a) Comparison of the TM-scores of the prediction models generated by VFlib and NNMake. (b) Comparison of the TM-scores of the prediction models generated by VFlib_random and NNMake_random. CD-HIT was used to remove all redundant proteins between the master library and benchmark proteins with pair-wise sequence identity > 30%.

**Table 1.**
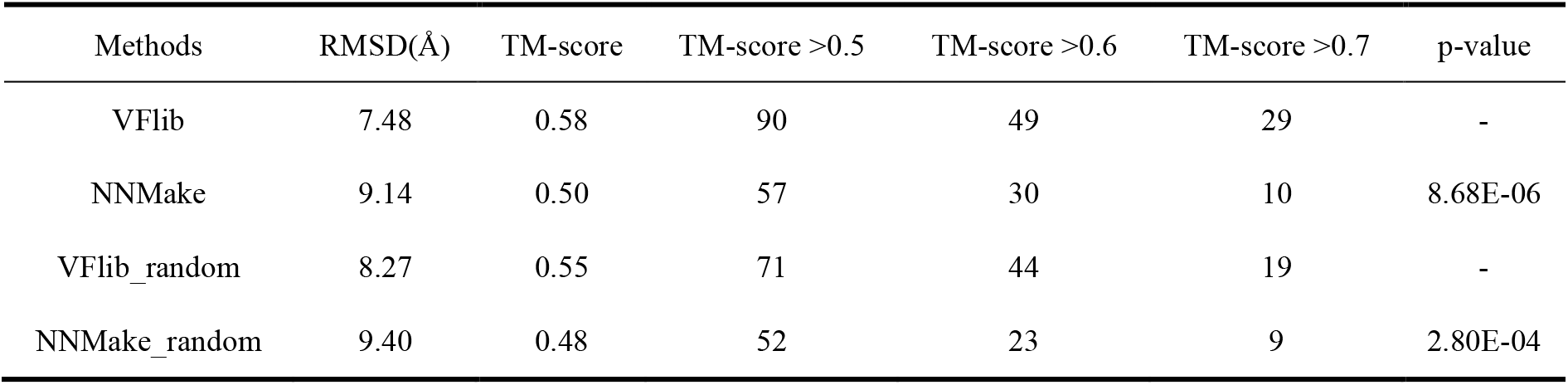
The results of VFlib, NNMake, VFlib_random and NNMake_random on 120 nonredundant proteins.

Since the strategy of assigning different probabilities to fragments based on distance information is designed for VFlib, the accuracy improvement effect of this strategy on the prediction model generated by using VFlib is more obvious than that of using NNMake. Therefore, we removed this strategy in the protein structure sampling program (VFlib_random and NNMake_random) to further verify the quality of the fragment libraries. The VFlib_random was tested on same 120 nonredundant benchmark proteins and compared with NNMake_random. Controlled experimental results in Table 1 illustrate that the average TM-score of the predicted models generated by VFlib_random (0.55) is 14.58% higher than that of NNMake_random (0.48), and the average RMSD of the predicted models generated by VFlib_random (8.27 Å) has reduced by 12.02% compared with that of NNMake_random (9.40 Å). Our method obtains 19 prediction models with TM-score >0.7, whereas NNMake_random obtains 9 prediction models with TM-score >0.7. Controlled experimental results in Figure 5 (b) show that VFlib_random achieves a higher TM-score than NNMake_random for 90 out of 120 nonredundant target proteins. The detailed results of TM-score and RMSD of the prediction models generated by VFlib, NNMake, VFlib_random and NNMake_random are shown in Supplementary Table S9.

The above controlled experimental results show that under the same assembly conditions, the variable-length (3-21) fragments generated by VFlib could generated a more accurate prediction model than NNMake. The fragments generated by the NNMake with fixed length and not diverse enough. This makes it difficult to form correct folds during the assembly process, especially for some β-sheet structures. Figure 6 shows the comparison of the native structure (protein ID: 4LE0_B) with the predicted models by VFlib and NNMake. The TM-score of VFlib (0.84) is significantly higher than that of NNMake (0.77). Although the prediction model obtained by NNMake forms a roughly correct topology, the native β-sheet structure region (gray) is erroneously formed as a loop structure (red). By contrast, the model obtained by VFlib has successfully formed the β-sheet structure (blue). This may be caused by the fixed length of the fragments generated by NNMake, which may destroy the complete secondary structure of the original protein. By contrast, the fragments generated by VFlib have variable lengths and enhanced diversity.

**Figure 6.**
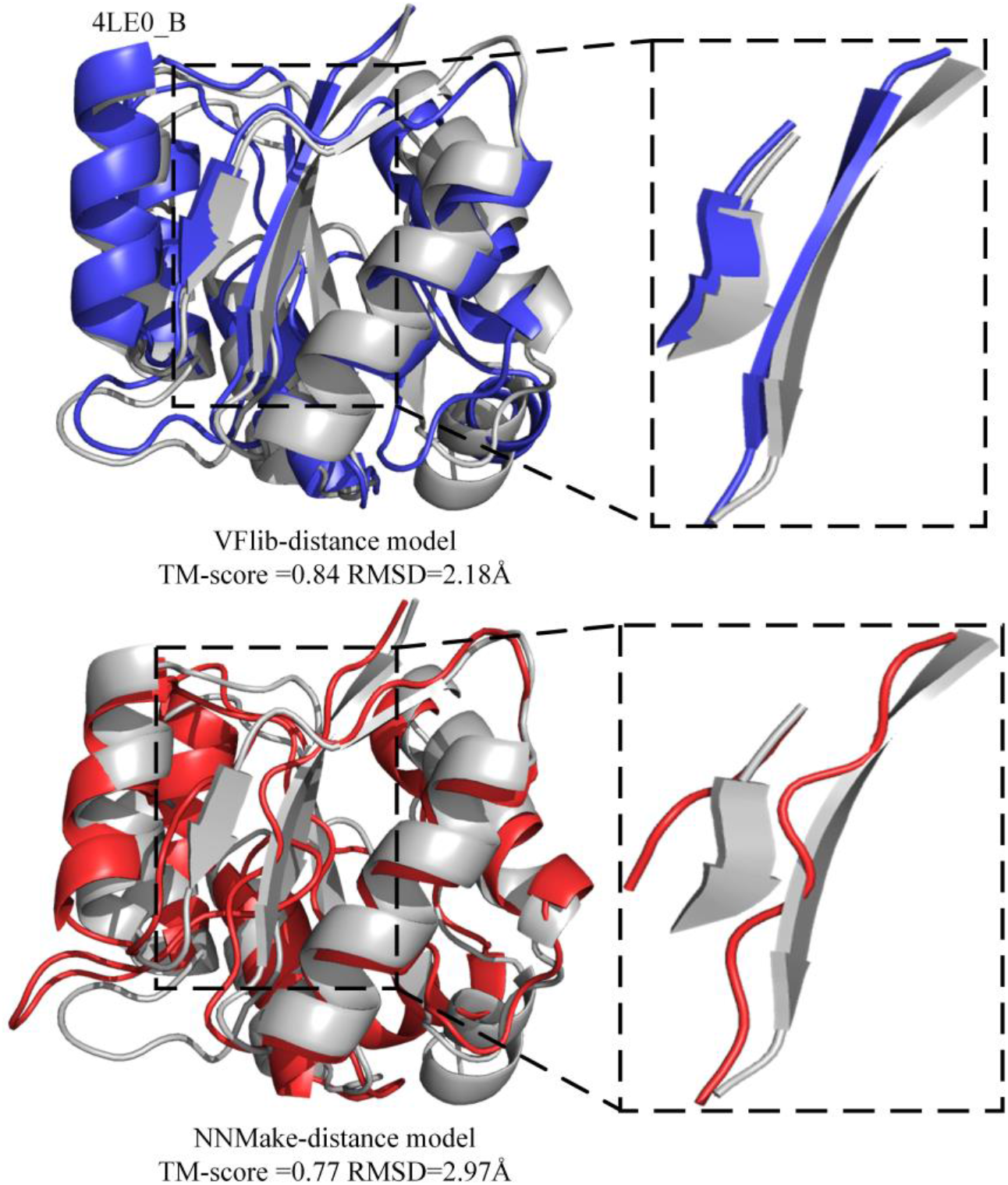
Comparison of the prediction models generated by VFlib (blue) and NNMake (red) with the native structure (gray) of the target protein (protein ID: 4LE0_B).

## 4. Conclusion

In this work, we construct a variable-length fragment library (VFlib) for *de novo* protein structure prediction. In VFlib, a master structure database was first constructed based on PDB (2021-08). Then, HMM-profile and secondary structure information of each protein in the master structure database was obtained by HHsuite and DSSP, respectively. In addition, a more complete dynamically variable-length profile-profile comparison strategy was proposed to make the fragment library of specific query sequence include more diverse fragments. The secondary structure information was used to further screen the retrieved fragments to generate the final variable-length fragment library.

In order to ensure the fairness of the experiment, it is necessary to remove homologues for avoiding the influence of the homologous protein in the master database. In 120 nonredundant benchmark proteins, the global precision and coverage of the fragment library generated by VFlib were 55.04% and 94.95% at the RMSD cutoff of 1.5 Å, respectively. Compared to the benchmark method of NNMake, VFlib have improvement in global precision with equivalent coverage. In addition, fragments generated by VFlib and NNMake were used to predict structure models based on Rosetta’s ClassicAbinitio protocol to further evaluate their quality. Controlled experimental results show that the accuracy of the prediction models generated by VFlib is greater than that of NNMake. The above experiments show that VFlib has higher quality and better diversity of fragments than NNMake. Notably, the proposed method relies on the diversity of the master database. As the number of experimentally determined proteins continues to increase, the fragment library generated by VFlib will be more diverse, which can not only better assist the fragment-based de novo prediction method, but it will also helpful the research of protein design [41–43].

## Key points

- We construct a variable-length fragment library for de novo protein structure prediction (VFlib), which is a valuable attempt to improve accuracy while ensuring high coverage through a more complete dynamically variable-length profile-profile comparison strategy and the further adjustments using secondary structure information.
- In VFlib, a master structure database was constructed based on PDB (2021-08), and the HMM-profile and secondary structure information of each protein in the master structure database was obtained by HHsuite and DSSP respectively.
- The experimental results on 120 nonredundant benchmark proteins show that the fragment library generated by our method has an improvement in the global precision with equivalent coverage compared to NNMake, and the performance of the predicted structure model using VFlib is also better.

## Supplementary Data

Supplementary data are available online at BIB.

## Data and code availability

All data needed to evaluate the conclusions are present in the paper and the Supplementary Materials. The additional data and code related to this paper can be downloaded from https://github.com/iobio-zjut/VFlib.

## Funding

This work has been supported in part by the National Nature Science Foundation of China (No. 62173304 and No. 61773346), the Key Project of Zhejiang Provincial Natural Science Foundation of China (No. LZ20F030002) and the National Key Research and Development Program of China (No. 2019YFE0126100).

## Notes

### Competing Interest Statement

The authors have declared no competing interest.

